# Zebrafish Rif1 impacts zygotic genome activation, replication timing, and sex determination

**DOI:** 10.1101/2022.10.21.512903

**Authors:** Emily A. Masser, Tyler D. Noble, Joseph C. Siefert, Duane Goins, Courtney G. Sansam, Christopher L. Sansam

## Abstract

Deregulated DNA replication causes human developmental disorders and cancer, but we know little about how DNA replication is coordinated with changes in transcription and chromatin structure. The initiation of replication forks follows a spatiotemporal pattern called the replication timing program. We have developed the zebrafish into a model system to study the mechanisms by which the replication timing program changes during the extensive changes in the cell cycle, transcription, chromatin organization, and nuclear structure that occur during development. Our previous studies identified changes in DNA replication timing patterns occurring from the onset of zygotic transcription through gastrulation in zebrafish embryos. Rif1 is required for DNA replication timing in a wide range of eukaryotes. The broader role of Rif1 in establishing the replication timing program and chromatin structure during early vertebrate development remains unknown. We have generated Rif1 mutant zebrafish and have performed RNA sequencing and whole-genome replication timing analyses on multiple developmental stages. Rif1 mutants were viable but had a defect in female sex determination. Surprisingly, Rif1 loss predominantly affected DNA replication timing after gastrulation, while its impacts on transcription were more substantial during zygotic genome activation. Our results indicate that Rif1 has distinct roles in DNA replication and transcription control that manifest at different stages of development.

## Introduction

The initiation of DNA replication follows a pattern where different genomic regions replicate at distinct times throughout S-phase. This temporal pattern is known as the replication-timing program (***Rhind and Gilbert, 2013***). Replication timing (RT) correlates with chromatin organization and gene activity (***Hiratani et al., 2010***). In multicellular organisms, early replication often correlates with active transcription; whereas, late replication frequently coincides with transcriptionally silent genes (***Hiratani et al., 2008***; ***MacAlpine et al., 2004***; ***Schübeler et al., 2002***; ***White et al., 2004***). In mammals, pluripotent cells exhibit a replication-timing pattern that changes with cellular differentiation; in addition, the switch in RT is related to changes in epigenetic modifications, transcriptional activity and the three-dimensional (3D) organization of chromatin (***Dixon et al., 2015***; ***Hiratani et al., 2010***, ***2008***; ***Rhind and Gilbert, 2013***; ***Zhou et al., 2002***). The causal relationships between DNA replication and developmental changes, such as transcription, chromatin structure and 3D chromatin organization have been unclear.

The Rif1 (Rap1interacting factor 1) protein is of particular interest because it has been implicated in the regulation of transcription, chromatin structure, 3D chromatin organization, DNA repair, and DNA replication timing. In fact, Rif1 is the single factor known to influence RT in all eukaryotes (***Cornacchia et al., 2012***; ***Hayano et al., 2012***; ***Hiraga et al., 2014***; ***Rhind and Gilbert, 2013***; ***Yamazaki et al., 2012***). Through conserved motifs, Rif1 has been shown to interact physically with the PP1 phosphatase (***Davé et al., 2014***; ***Hiraga et al., 2014***; ***Mattarocci et al., 2014***; ***Sreesankar et al., 2015***; ***Sukackaite et al., 2017***). Rif1 controls RT by bringing PP1 to latereplicating regions of the genome, thereby causing de-phosphorylation of DNA replication factors during early S-phase and delaying initiation (***Davé et al., 2014***; ***Foti et al., 2016***; ***Hiraga et al., 2014***, ***2017***; ***Mattarocci et al., 2014***; ***Sreesankar et al., 2015***; ***Yamazaki et al., 2012***).

Our understanding of how Rif1 affects RT during embryonic development is incomplete. RT is developmentally regulated through Rif1 in *D. melanogaster* (***Seller and O’Farrell, 2018***). In early fly embryos, Rif1-activation initiates late replication of satellite repeats and causes the first lengthening of the cell cycle (***Seller and O’Farrell, 2018***). In Xenopus laevis, Rif1 knockdown accelerates early embryonic cell cycles and disrupts RT patterns in retinal progenitor cells (***Meléndez García et al., 2022***). Loss of Rif1 in mice is embryonic lethal with partial penetrance (***Buonomo et al., 2009***; ***Chapman et al., 2013***; ***Daxinger et al., 2013***; ***Enervald et al., 2021***). Cell lines derived from Rif1 mutant mice display altered RT profiles, but the extent to which RT is disrupted in the developing embryo has not been shown. Although Rif1 loss causes RT changes in cell lines from a wide range of species, it is unknown whether Rif1 plays roles in the establishment or change of the RT program during vertebrate embryogenesis.

We have developed the zebrafish into a model system to study how modifications in RT coincide with extensive changes in cell cycle and transcription that occur in the developing embryo. The earliest stages of zebrafish development have been extensively studied, so it is possible to collect large numbers of synchronously developing embryos at developmental time points when cells are known to be pluripotent, specified or committed to particular cell fates (***Ho and Kimmel, 1993***). As a result, it is feasible to study how changes in RT accompany developmentally regulated changes during cellular differentiation in vertebrates. We previously mapped RT at various stages of early wild-type zebrafish development (***Siefert et al., 2017***). We revealed that a weak RT pattern present at the onset of zygotic transcription becomes progressively more defined as the fish develop. In addition, during gastrulation, large genomic domains switch from early-to-late replication in the embryo. Finally, we observed that changes in epigenetic modifications involved in enhancer activity correlate with developmental changes in RT (***Siefert et al., 2017***).

To reveal the function of Rif1 in gene expression and RT during vertebrate development, we have generated Rif1 mutant zebrafish, and we have performed RNA sequencing and whole-genome RT analyses before the first major wave of zygotic transcription and before and after gastrulation. We found that Rif1 is not essential for embryonic development, but its loss impairs female sex de-termination. Although the Rif1 mutants had altered RT at each stage, the effects of Rif1 loss on RT were minor compared to changes that naturally occur during development. DNA replication timing was most affected by Rif1 loss after gastrulation, while transcription was most affected early in development. Rif1 was required for normal expression of genes during the first wave of zygotic genome activation. Our results indicate that in zebrafish, Rif1 has distinct roles in DNA replication and transcription control that manifest at different stages of development.

## Results

### Generation of a Rif1 loss zebrafish model

The zebrafish genome has a single copy of the *rif1* gene per haploid genome on chromosome 9 (GRCz11 22,780,901-22,802,172). Similar to other vertebrates, the *zrif1* gene encodes a 2,347 amino acid protein with heat repeats and PP1 phosphatase interaction sequences (***Figure 1***A,B) **(*Alavi et al., 2021*)**. To determine the effects of Rif1 loss-of-function in the zebrafish, we mutated the gene using a transcription activator-like effector nuclease targeting exon 8 (TALEN; ***Figure 1***A). Frameshift mutations in exon eight would likely induce nonsense-mediated decay (NMD) of the mutant mRNA. If expressed, the mutant allele would produce a protein ending within the sequence encoding the heat repeats (***Figure 1***A). We selected a stable line **(*rif1omf201*)** with a seven base-pair frameshift mutation for further analysis (***Figure 1***A).

**Figure 1.**
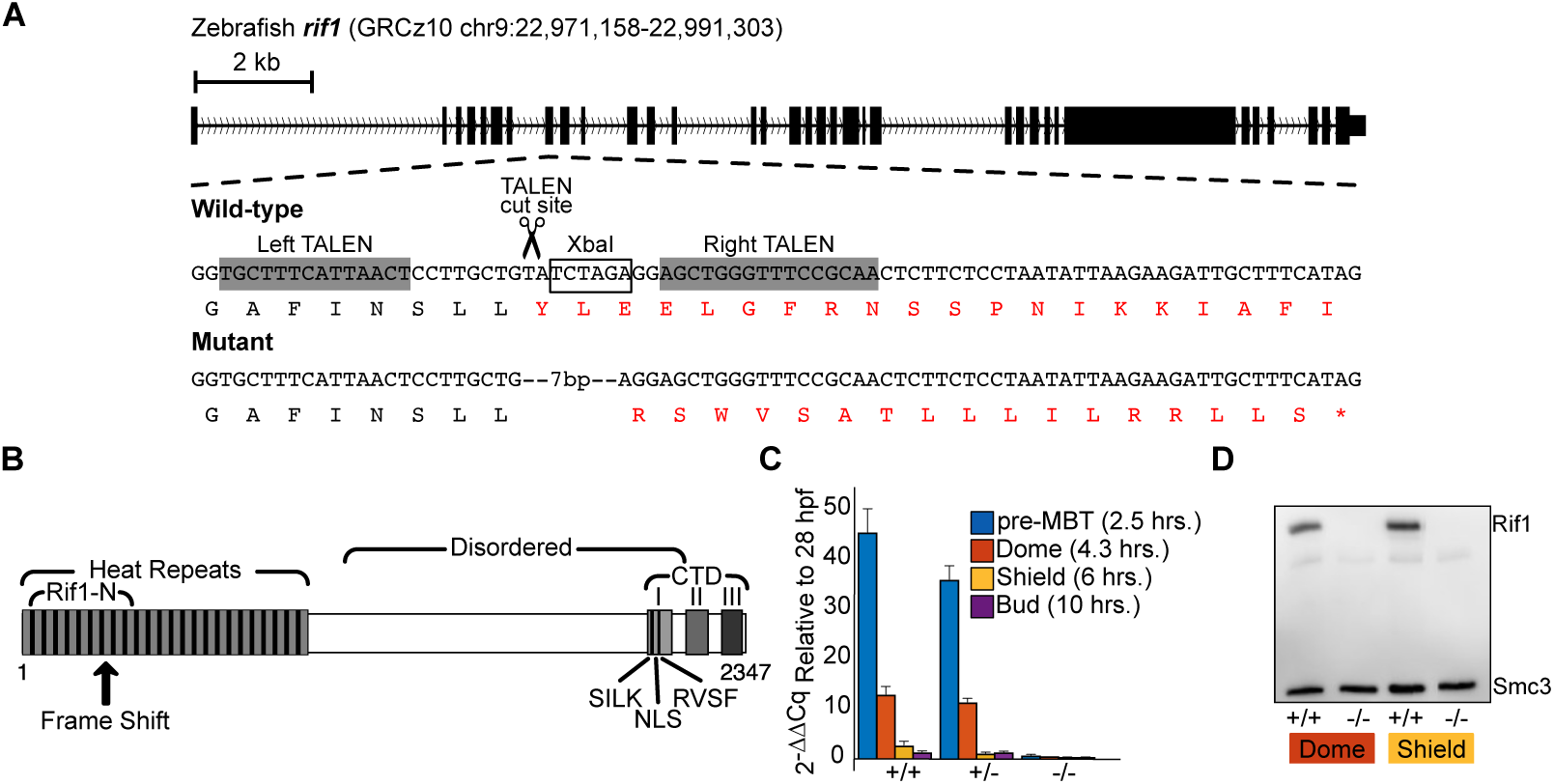
A TALEN-induced mutation abrogates ***rif1*** expression in zebrafish. (A) Schematic illustration of the TALEN mutation in exon 8 of the zebrafish *rif1* gene. The DNA and protein sequences at the TALEN site are shown for the wild-type and a 7 bp deletion mutant. The deletion causes a frame shift and introduces a premature termination codon. The affected amino acids are in red. (B) Diagram of the Rif1 protein. The 7 bp frame shift mutation occurs in the conserved Rif1 N-terminal domain (Rif1-N), causing truncation of the Heat repeats and loss of the nuclear localization signal (NLS) and the PP1 phosphatase SILK and RVSF interaction motifs of the C-terminal domain (CTD). (C) Expression levels of *rif1* measured by quantitative RT-PCR analysis at pre-MBT (2.5 hours), dome (4.3 hours), shield (6 hours) and bud (10 hours) in wild-type (+/+), heterozygous (+/-) and homozygous (-/-) *rif1* zebrafish. Error bars represent standard deviation of the mean. (D) Rif1 and Smc3 protein expression levels were analyzed by immunoblotting at dome and shield in wild-type (+/+) and homozygous (-/-) *rif1* zebrafish.

We first tested whether zygotic *rif1omf201* mutants developed normally and survived to sexual maturity. Most offspring from *rif1omf201/+* crosses hatched and began free feeding by five days post-fertilization (dpf; data not shown). Therefore, we generated clutches of *rif1+/+*, *rif1omf201/+*, or *rif1omf201/omf201* embryos spawning F2 mutant and wild-type fish. We then collected embryos of each genotype at 2.5, 4.3, 6, and 10 hours post-fertilization (hpf) and measured *rif1* mRNA by quantitative RT-PCR (***Figure 1***C). These developmental time points span the start of zygotic transcription through gastrulation **(*Kimmel et al., 1995*)**. Our data revealed that *rif1* expression in wild-type zebrafish embryos is developmentally regulated, with the highest expression level at 2.5 hpf and then a precipitous drop at all stages after that (***Figure 1***C). The significance of this change in mRNA expression for development is unknown, but we also found that Rif1 protein also decreases in the embryo over these stages (data not shown). The expression level of *rif1* mRNA in heterozygous and homozygous mutant fish was lower than wild-types at all stages (***Figure 1***C). In maternal-zygotic homozygous mutants, *rif1* mRNA was nearly undetectable (***Figure 1***C). Therefore, we conclude the mutation causes a substantial decrease in *rif1* mRNA.

To determine if Rif1 protein levels were also affected by the mutation, we generated a rabbit polyclonal antibody against the N-terminus of the protein (amino acids 1-150). The antibody’s antigen included the portion of the truncated protein that the mutant allele would express (***Figure 1***A, B). We did not detect any residual full-length Rif1 protein on immunoblots of nuclear lysates from maternal-zygotic homozygous mutants (***Figure 1***D). Furthermore, we have not found evidence on the immunoblots that the *rif1omf201* mutants express truncated Rif1 protein (***Figure 1***D). Altogether, these data suggest that the *rif1omf201* mutation eliminates Rif1 protein expression in the embryo.

### Rif1 is not essential in zebrafish but its loss affects early development and sex determination

Given that Rif1 loss in mice caused incompletely penetrant embryonic lethality, we decided to track the development and survival of a large cohort of *rif1omf201* mutants **(*Buonomo et al., 2009*)**. We raised 766 offspring from *rif1omf201/+* in-crosses and counted fish of each genotype when they reached sexual maturity (***Figure*** 2A, B). Zebrafish sex is determined by poorly defined genetic and environmental factors, so we also counted fish of each gender **(*Kossack and Draper, 2019*)**. Although all genotypes were at their expected Mendelian proportions, the number of female heterozygous and homozygous mutants was reduced (***Figure*** 2B, C). We conclude that although the *rif1omf201* mutation is not zygotic lethal, it does perturb sex determination.

**Figure 2.**
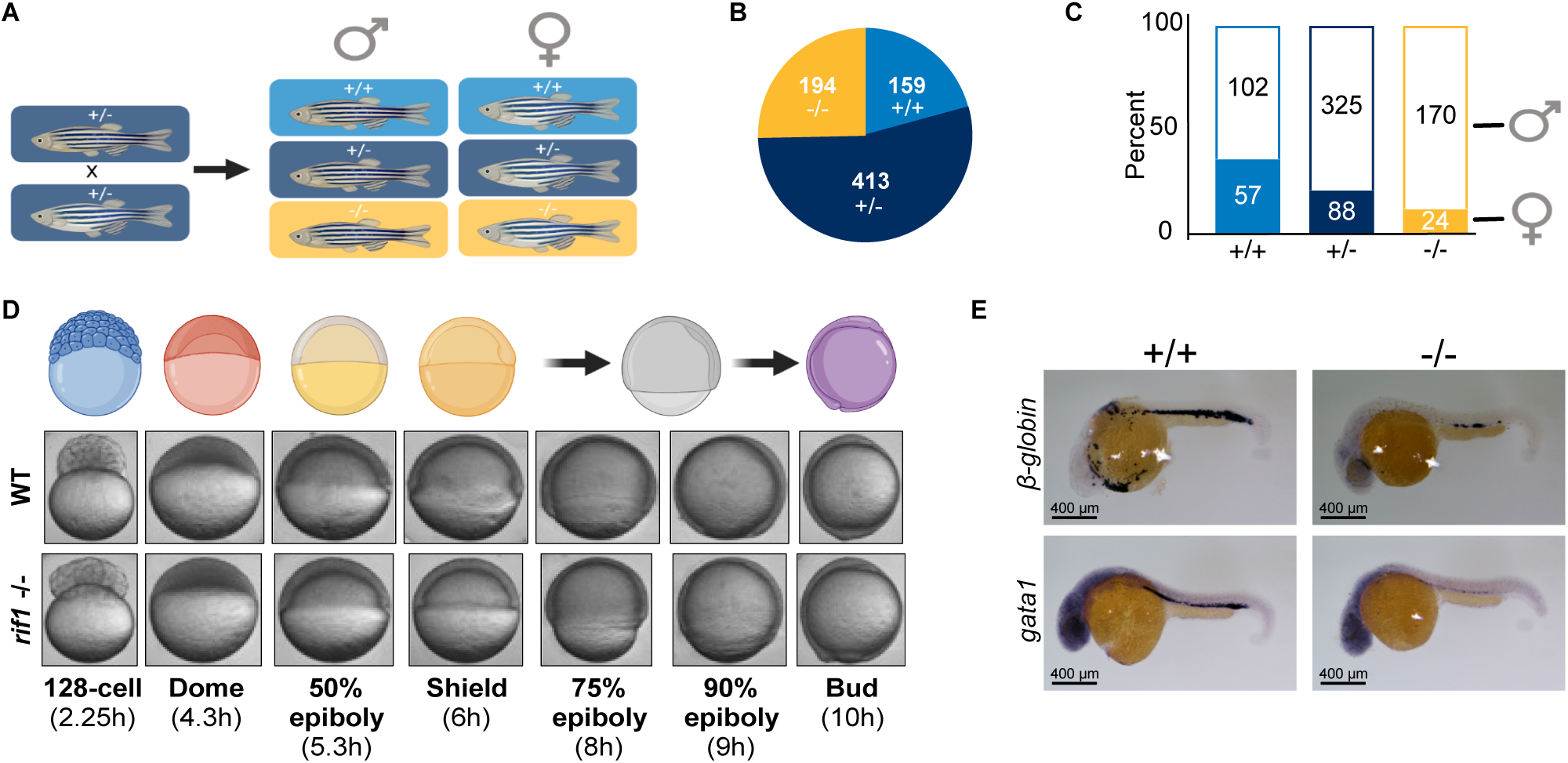
Loss of ***rif1*** affects zebrafish sex determination, epiboly, and primitive erythropoiesis. (A) Schematic of the heterozygous *rif1* mutant crosses used to test the essentiality of *rif1*. 766 fish were raised to sexual maturity (3 months) and then genotyped. (B) Quantitation of the genotypes from the heterozygous crosses shows the expected Mendelian ratios. (C) Counts of males and females from the heterozygous crosses shows fewer than expected *rif1* mutant female fish. (D) Time-lapse imaging of wild-type and maternal-zygotic *rif1* -/-early development shows a delay in epiboly completion in the mutants. (E) Expression of gata1 and B-globin at 24 hpf marks primitive erythroid cells. In situ hybridization shows that gata1 and B-globin expression at 24 hpf was decreased in *rif1* homozygous mutants, indicating loss of primitive erythroid cells.

Our qRT-PCR analysis suggested that zebrafish have substantial *rif1* mRNA at pre-MBT (midblastula transition), before full zygotic genome activation (***Figure 1***A). Maternally supplied *rif1* mRNA or protein might mask critical functions during early development in zygotic mutants. Therefore, we carefully analyzed the maternal-zygotic *rif1omf201/omf201* mutants, which lack maternally supplied *rif1*, for any overt morphological defects during development. Mutant fish development was delayed by approximately 1 hour at the end of gastrulation, as indicated by slower epiboly completion (10 hpf; ***Figure 2***D). During epiboly, the blastoderm and yolk syncytial layer move toward the vegetal pole to envelop the yolk cell. From the dome stage onward, gastrulation stages are commonly designated by the percentage of the yolk cell covered by the blastoderm. The delay in *rif1omf201/omf201* mutants was evident as slower progression through the 75% and 90% epiboly stages (***Figure 2***D). To quantify this phenotype, we scored the number of embryos that had completed epiboly, defined as 100% yolk coverage, at 10 hpf. Most wild-type embryos had completed epiboly by 10 hpf (18 of 24), whereas only 2 of 24 mutant embryos had completed epiboly at this time. By 11 hpf, all wild-type and mutant embryos had completed epiboly. Moreover, *rif1omf201/omf201* mutants had fewer circulating blood cells relative to wild types at one dpf. Consistent with the blood cell deficiency, *rif1omf201/omf201* mutants expressed lower levels of hematopoiesis markers *β*-*globin* and *gata-1*; (***Figure 2***E). Despite the delay in early development and the deficiency in primitive hematopoiesis, maternal-zygotic *rif1omf201/omf201* mutants recover and develop to adulthood.

### Rif1 loss impairs the establishment of the RT program genome-wide during zebrafish development

Given that late replication and S-phase lengthening requires Rif1 in Drosophila, we hypothesized that RT program establishment and change in vertebrates would also depend on Rif1 **(*Seller and O’Farrell, 2018*)**. We tested this hypothesis by profiling RT in wild-type and *rif1omf201/omf201* mutant zebrafish embryos at multiple stages. We focused on early development because we had previously described changes in RT at multiple time points from the 256-cell through gastrulation in zebrafish **(*Siefert et al., 2017*)**. We showed that the 256-cell and shield stages in zebrafish are when the RT pattern shifts from being relatively weak to defined **(*Siefert et al., 2017*)**. Furthermore, the major wave of zygotic genome activation occurs around the 256-cell stage. Therefore, 256-to-shield is analogous to the stages in Drosophila development when Rif1 promotes the onset of late replication of satellite repeats **(*Seller and O’Farrell, 2018*)**. We also previously observed that the shield and bud stages are when numerous broad RT changes occur **(*Siefert et al., 2017*)**. Therefore, we used whole-genome sequencing to generate RT maps for wild-type and Rif1 mutant zebrafish embryos at 128-cell/pre-MBT (2.25 hours), shield (6 hours), bud (10 hours), and 24 hpf. Replication timing was inferred from relative copy number in S-phase and G1-phase DNA. Regions that replicate early are overrepresented in S-phase DNA relative to G1 DNA, whereas regions that replicate later show lower S/G1 enrichment.

We first assessed overall differences in RT between all genotypes and stages with hierarchical clustering on Euclidean distances between samples (***Figure 3***A). Using this approach, we found that samples clustered first by developmental stage and then genotype, demonstrating that the RT changes that occur during zebrafish development are more significant than those caused by Rif1 loss. The Euclidean distances between wild-type and *rif1omf201/omf201* samples were minor at the pre-MBT stage and increased progressively through development, reaching their highest levels at 24 hpf. Importantly, biological replicates of the same developmental stage and genotype were most similar, demonstrating the reproducibility of our RT measurements.

**Figure 3.**
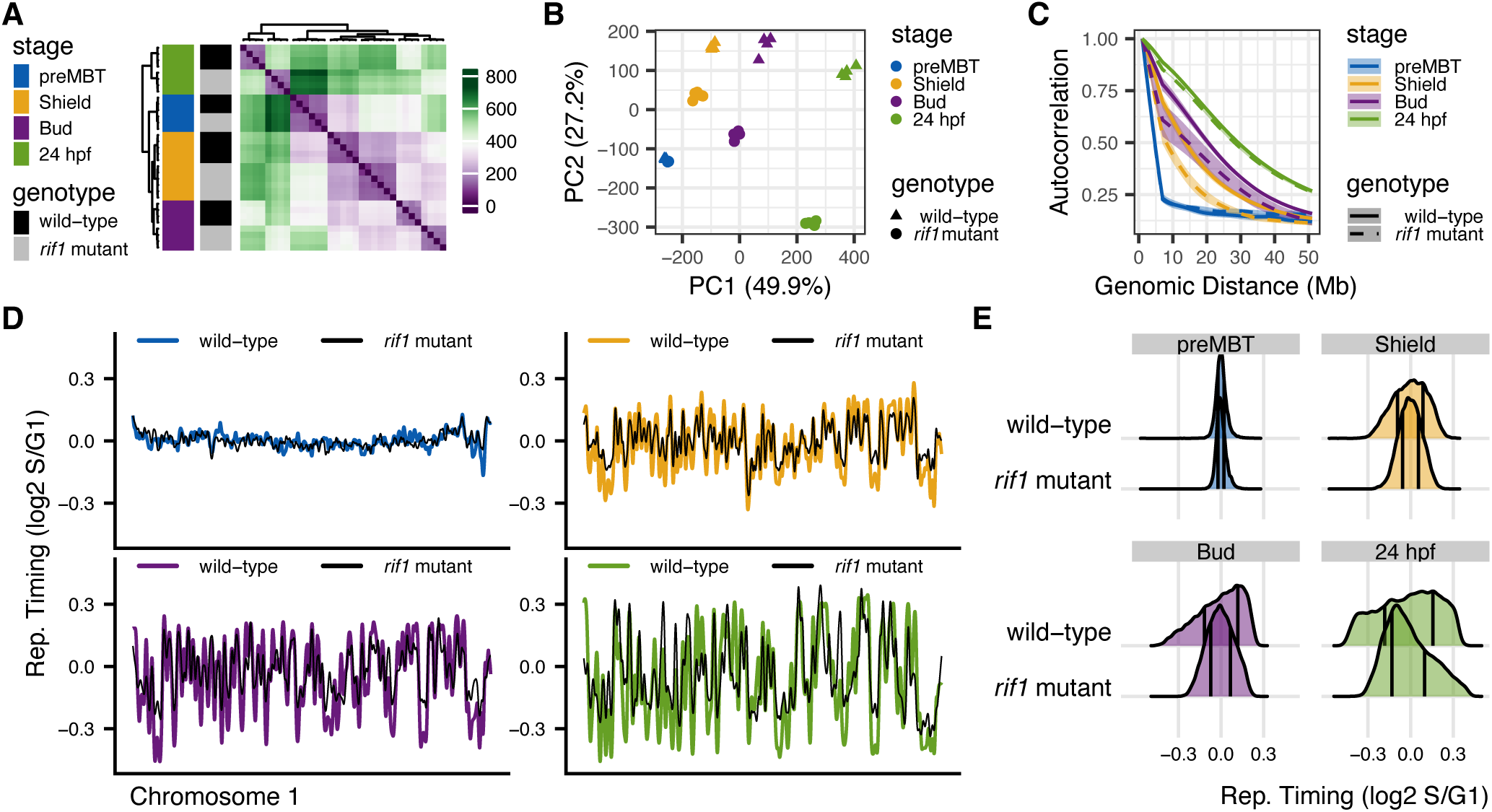
Rif1 loss reduces the general sharpening of RT patterns that occur during normal zebrafish development. (A) Heat plot and dendograms showing hierarchical clustering of samples based on Euclidean distances calculated with RT values. (B) Principle component analysis reveals that 50% and 27% of variance in RT between samples is due to developmental stage (PC1) and genotype (PC2), respectively. (C) Autocorrelation analysis of RT values shows that RT profiles gradually gain structure during zebrafish development, and Rif1 loss delays that process. (D) Whole chromosome 1 RT profiles are shown for preMBT (top left), shield (top right), bud (bottom left) and 24 hpf (bottom right) stages. The RT profiles gradually develop definition, but Rif1 loss reduces the amplitude of the peaks and valleys. (E) Density plots of genome-wide RT values show a reduced dispersion of values from 0 in the *rif1* mutants compared to wild-types. Vertical black lines within density plots represent 25 th and 75 th percentiles.

The conclusions drawn from the PCA were the same as Euclidean distance analysis (***Figure 3***B). All biological replicates clustered together on the PC2 vs. PC1 plot, showing that the data is reproducible. Principal components 1 and 2 described RT differences caused by developmental stage and genotype, respectively. The first principal component, which explained 50% of the RT variance, separated samples by developmental stage; the second component, which explained 27% of the variance, separated samples by genotype. Therefore, the developmental stage affected RT more strongly than the genotype. There was no difference between wild-type and *rif1omf201/omf201* samples in the first two principal components at the pre-MBT stage (***Figure 3***B; blue points), supporting that Rif1-loss had little effect on RT early in development. As development proceeded, wild-type and *rif1omf201/omf201* mutant samples progressively separated on the second principal component, supporting that RT changes caused by Rif1 loss were more pronounced later in development.

We previously showed that the RT pattern in zebrafish embryos gradually gained the hallmark peak-and-valley RT profile structure as development proceeded **(*Siefert et al., 2017*)**. To quantify this structure, we used autocorrelation analysis, which measures how similar RT values are across increasing genomic distances along the same chromosome. Higher autocorrelation over hundreds of kilobases indicates a more continuous domain-scale RT profile. We calculated the autocorrelation of RT values along each chromosome in wild-type and *rif1omf201/omf201* embryos at each developmental stage (***Figure 3***C). As described previously, RT value autocorrelation increased between each stage in wild-type embryos **(*Siefert et al., 2017*)**, indicating that the timing program progressively gained domain-scale structure during development. The autocorrelation of *rif1omf201/omf201* RT values also increased during development, but this change lagged behind wild-type. For example, the autocorrelation of *rif1omf201/omf201* RT values at the bud stage resembled that of wild-type at shield. The autocorrelation analysis indicated that Rif1 loss delayed the gradual gain in RT profile structure during development.

Broad differences in RT were visually evident from plots of normalized (log2 S/G1) RT values at all stages after the MBT (***Figure 3***D). At the 128-cell stage (before the MBT), both *rif1omf201/omf201* and wild-type RT profiles were flat compared to those at later stages (***Figure 3***D). This was consistent with replication origin firing synchrony at the 128-cell stage in both the wild-type and *rif1* mutants. After the 128-cell stage, the RT profiles of *rif1omf201/omf201* and wild-type embryos developed peak-and-valley structures, but the *rif1omf201/omf201* profiles were less defined (***Figure 3***D; compare black lines to colored lines). At each stage after 128-cell, the amplitudes of peaks and valleys appeared lower in the *rif1omf201/omf201* profiles (***Figure 3***D). Overall, the RT values (log2 S/G1) in *rif1omf201/omf201* mutants at shield, bud, or 24 hpf were less dispersed from the median than the RT values of wild-type embryos at the same stage (***Figure 3***E). Altogether our data show that Rif1 sharpens the genome-wide RT profile during zebrafish development.

### Rif1 is not required for major changes in RT that occur during zebrafish development

We previously identified two significant changes in RT that occur during early zebrafish development **(*Siefert et al., 2017*)**. First, genomic domains spanning hundreds of kilobases underwent late-to-early changes in RT. Both coincided with H3K27 acetylation or DNA demethylation at putative enhancers within the domains. Second, ∼40 million base-pairs of zebrafish chromosome 4q and a large chromosomal domain on chromosome 22 (5,749,848-10,129,684) both underwent early-to-late RT switches during gastrulation. The mechanisms underlying those changes are unknown.

To show whether RT changes that coincide with enhancer activation require Rif1, we calculated RT values at putative enhancers that become acetylated during early zebrafish development (***Figure 4*A-B**) **(*Bogdanovic et al., 2012*)**. The majority (1,768/2,498) of differentially acetylated regions (DARs) that become acetylated between 4.3–24 hpf changed to earlier replicating between the shield (6 hpf) and 24 hpf stages in both wild-type and *rif1omf201/omf201* mutant embryos. Moreover, the shield-to-24 hpf changes in RT for wild-type and *rif1omf201/omf201* DARs were strongly correlated (rs=0.76, p<2.2e-16; ***Figure 4*A**). Therefore, we conclude that Rif1-loss does not generally affect the changes in RT that coincide with enhancer activation during early zebrafish development.

**Figure 4.**
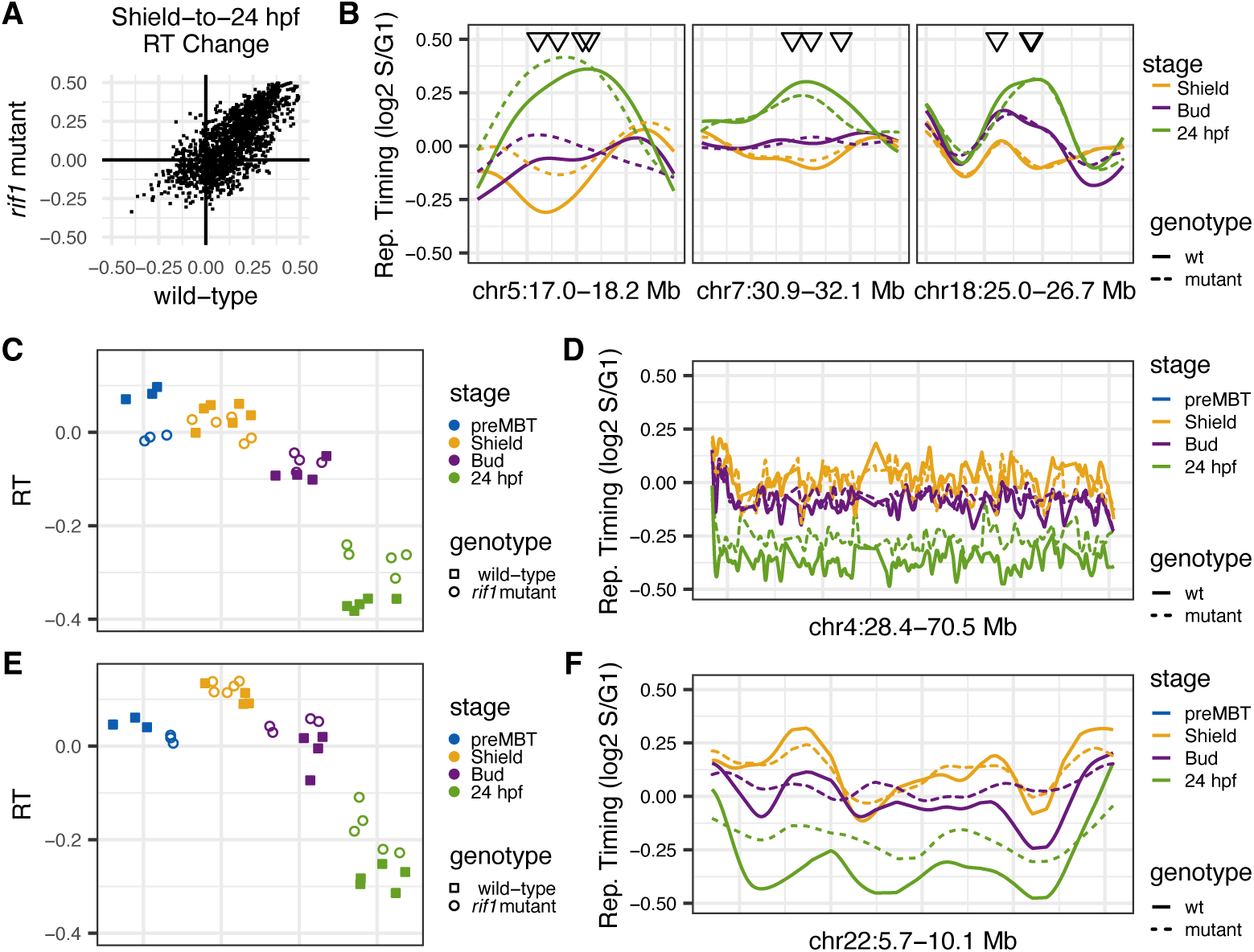
Rif1 loss does not prevent major changes in RT that occur during early zebrafish development. (A) Scatterplot of changes in RT of 2,498 putative enhancers that are H3K27 acetylated during gastrulation (DARs). A positive shield-to-24 hpf change indicates a switch to earlier replication during gastrulation or segmentation. The majority of acetylated regions switch to earlier replication in both wild-type and *rif1* mutants. (B) RT plotted across three regions with representative DARs. The positions of individual DARs are marked with triangles. (C-F) RT calculated for all of chr4 : 28,400,000-70,500,000 (C-D) or chr22 : 5,700,000-10,100,000 (E-F) at each stage and genotype. In C and E, point shape denotes genotype, with open squares representing wild-type samples and open circles representing *rif1* mutant samples; point color denotes developmental stage. Panels B, D, and F show shield, bud, and 24 hpf data; pre-MBT values are not plotted in these panels. The broad early-to-late switches in RT during gastrulation occur in the *rif1* mutants.

To determine whether the early-to-late switches in RT that occur on chromosomes 4 and 22 require Rif1, we calculated RT values for Chr4q or Chr22:5,749,848-10,129,684 for both wild-type and *rif1omf201/omf201* embryos at 128-cell, shield, bud, and 24 hpf stages (***Figure 4*C-F**). Unexpectedly, Chr4q was replicated earlier during S-phase in the *rif1omf201/omf201* embryos than in wild-type embryos at the 128-cell stage, but that difference was minor (***Figure*** 4C, D). As we previously described, early-to-late RT switches on Chr4q (ANOVA p=7.27e-05) and Chr22 (ANOVA p=7.27e-05) occurred between the shield and 24 hpf stages in wild-type embryos (***Figure 4***C-E). The early-to-late switches at Chr4q (ANOVA p=0.803) and Chr22 (ANOVA p=0.803) also occurred in the *rif1omf201/omf201* mutant embryos (***Figure 4***C-E). Although both Chr4q and Chr22:5,749,848-10,129,684 underwent early-to-late switches in the *rif1omf201/omf201* mutant embryos, both were earlier replicating in the mutant compared to wild-types at 24 hpf (***Figure 4***C-F). These differences might indicate that those specific timing switches were delayed or incomplete, or it could be due to the genome-wide flattening of the RT profiles. Indeed, when we compared genome-wide replication timing values from all genomic windows in wild-type and *rif1omf201/omf201* mutant embryos at 24 hpf, we observed that most late-replicating regions were earlier replicating in *rif1omf201/omf201* mutants (***Figure*** 3D, E). These data have led us to conclude that although Rif1 promotes the establishment of the RT profile genomewide, it does not generally affect specific RT switches during zebrafish development.

### Rif1 affects transcription at the earliest stages of zebrafish development

Mouse studies have revealed roles for Rif1 in regulating gene expression and epigenetic marks in embryonic stem cells and cancer cell lines **(*Dan et al., 2014***; ***Foti et al., 2016***; ***Klein et al., 2021***). It has been suggested that Rif1 affects transcription indirectly, possibly through its RT function **(*Klein et al., 2021*)**. We were particularly interested in whether Rif1 loss would cause transcriptional changes later in development when the RT program is established or in the early embryo, when there is a minimal RT pattern. Therefore, we used 3’ end RNA sequencing on maternal-zygotic *rif1omf201/omf201* mutant and wild-type zebrafish embryos at 1-cell (0.2 hours), 256-cell (2.5 hours), dome (4.3 hours), shield (6 hours), bud (10 hours) and day 1 (28 hours) (***Figure 5***).

**Figure 5.**
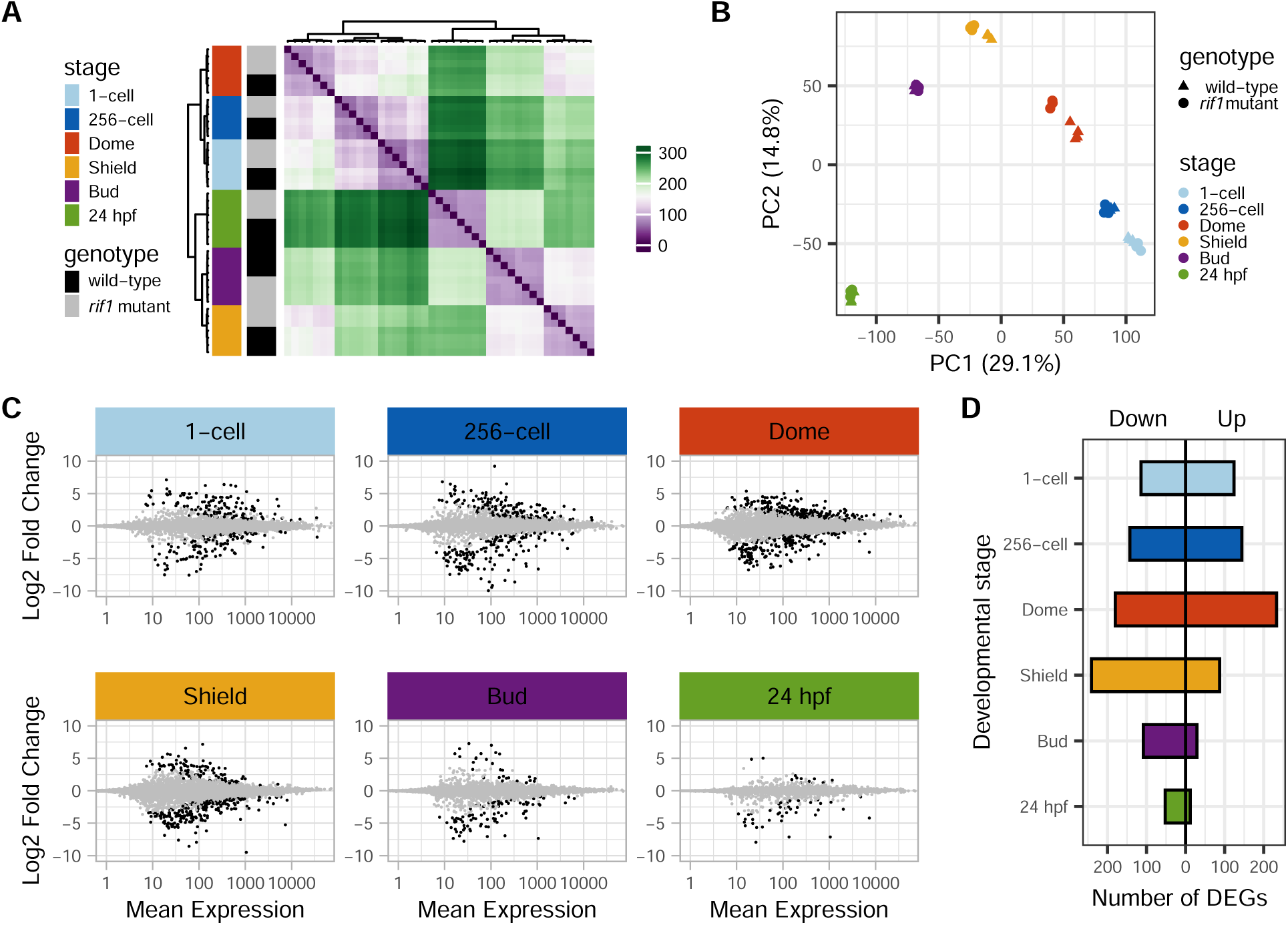
Rif1 loss affects gene expression preferentially during the earliest stages of zebrafish development. (A) Matrix with colors and dendrograms representing Euclidean distances between 3’ mRNA-Seq samples. The order of samples is based on hierarchical clustering and is the same for rows and columns. Each sample’s genotype and developmental stage is marked by a color to the left of its row. (B) Principal component analysis (PCA) plot showing clusters of samples based on similarities of 3’ mRNA-Seq data. The first two principal components are plotted, which represent 29.1% and 14.8% of the variance. Both the Euclidean distance and PCA analyses show that biological replicates are most similar, followed by samples from the same genotype and then stage. Rif1 mutant and wild-type gene expression differs most at dome and shield stages. (C) Fold-change in expression between *rif1* -/-and wild-type embryos versus normalized mean 3’ mRNA-Seq read counts. Black data points represent differentially expressed genes (DEGs) with statistically significant (adjusted Wald test p-value < 0.05; Log2FC > 0.5) changes in expression in the *rif1* mutants. (D) Counts of DEGs from C with lower (Down) or higher (Up) expression in *rif1* mutants.

We determined the overall effects of Rif1 loss on the embryonic transcriptome by analyzing the RNAseq data using principle component analysis (PCA) and hierarchical clustering based on Euclidean distances (***Figure*** 5A, B). We found that biological replicates were highly similar, and as expected, the transcriptome changed significantly between developmental stages. Rif1 loss impacted the transcriptome at every stage evaluated but not all stages were equally affected, with the most significant effects on gene expression at the dome and shield stages (***Figure 5***B). From this point forward, we will refer to differentially expressed genes (DEGs) with higher or lower expression

in the maternal-zygotic *rif1omf201/omf201* mutants as Up-DEGs or Down-DEGs, respectively (adjusted Wald test p value < 0.05 and Log2-fold change > 0.5 or < -0.5). The number of Up-DEGs peaked at the dome stage, while the number of Down-DEGs peaked at the shield stage (***Figure 5***C-D). To test whether transcriptionally affected genes were associated with replication timing state, we assigned each gene the nearest smoothed replication timing value and compared genes with increased, decreased, or unchanged transcript abundance at Dome. Genes with altered transcript abundance at Dome were not obviously enriched for early- or late-replicating regions relative to genes with no significant transcript change (Figure 5–figure supplement 2).

Zebrafish contribute most zygotic transcripts maternally, so the differences in transcript abundance in the *rif1omf201/omf201* mutants could be due to changes in the supply or stability of maternal mRNA **(*Harvey et al., 2013*)**. We used k-means clustering to categorize all DEGs based on their log-fold change in expression across all stages (Figure 5–figure supplement 1A, B). The majority of one-cell DEGs are expressed normally at other stages, and the majority of DEGs at other stages are expressed normally at one-cell (Figure 5–figure supplement 1A, B). Therefore, the differential expression of mRNAs at later stages in the *rif1omf201/omf201* mutants is not due to differential maternal contribution.

### Rif1 broadly suppresses transcription during Zygotic Genome Activation

To learn more about the genes affected by Rif1 loss, we looked at when zebrafish embryos normally express the DEGs. White et al. profiled the zebrafish transcriptome at 18 developmental time points **(*White et al., 2017*)**. We used this data set to categorize dome DEGs based on when their peak expression occurs in wild-type embryos (***Figure 6***A-D). We focused on dome DEGs because they were the most abundant. Dome Up-DEGs are enriched in the Blastula/Gastrula expression cluster (***Figure*** 6A, C). Since mRNAs that peak at the Blastula and Gastrula stages are likely to be transcribed during the major wave of zygotic genome activation at the MBT, our data suggests Rif1 loss stimulates the transcription of mRNAs at the first major wave of Zygotic Genome Activation (ZGA). Dome Down-DEGs are enriched in the pre-ZGA cluster, which suggests that Rif1 loss also enhances the destruction of maternal mRNAs (***Figure 6***B, D). The degradation of many maternal mRNAs in zebrafish is triggered by transcription of the *mir-430* microRNA cluster before the major wave of zygotic transcription **(*Giraldez et al., 2006*)**.

**Figure 6.**
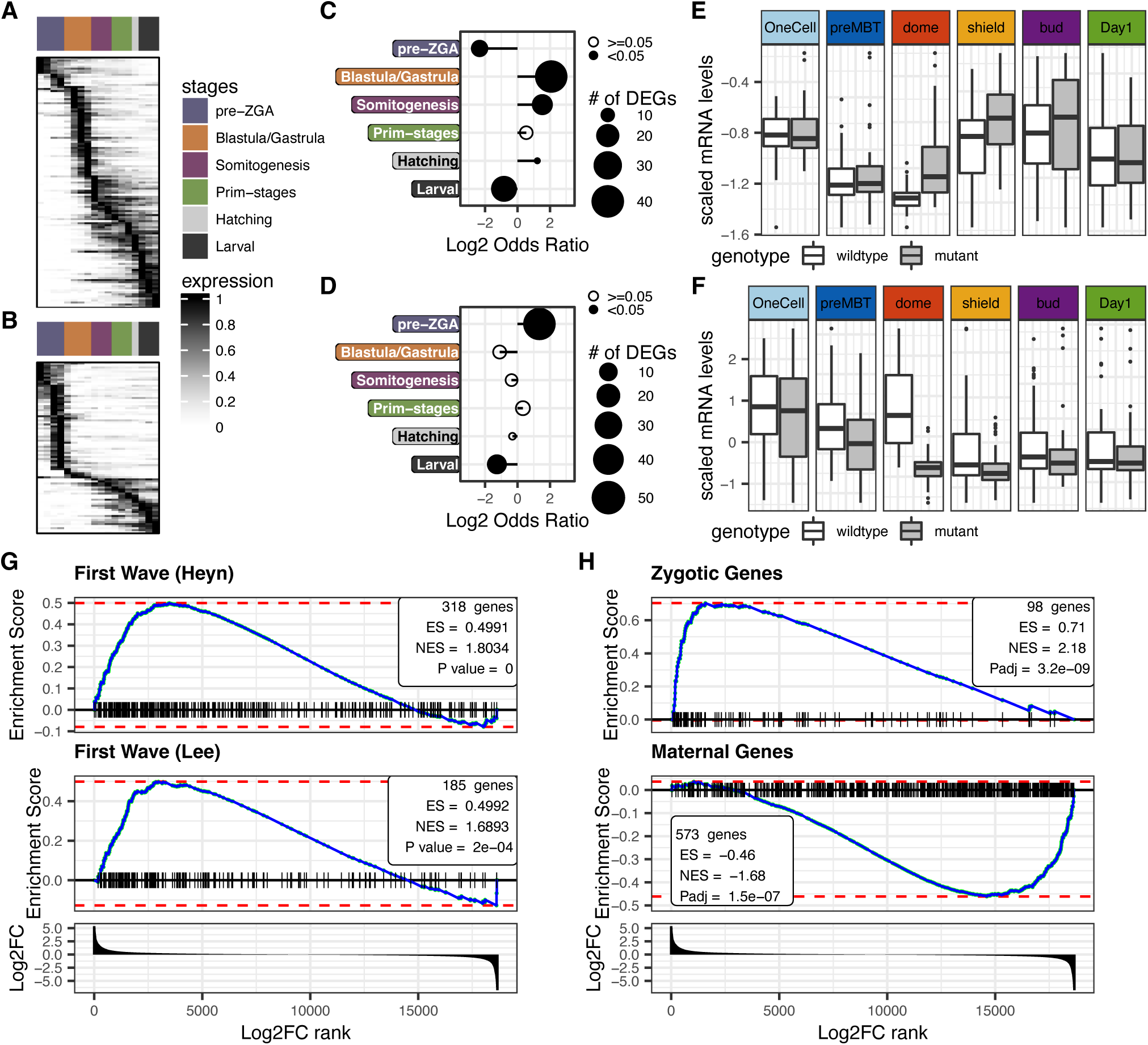
ZGA transcribed or degraded mRNAs are deregulated in ***rif1*** mutants. (A-B) Heat plots showing the expression (tags per million scaled to maximum) of *rif1* mutant Dome Up (A) or Down (B) DEGs (rows) across 18 developmental time points (columns) in wild type embryos. Expression data is from White, et al. **(*White et al., 2017*)**. The legend across the top of each heat plot denotes six developmental phases. (C-D) Log2 odds ratios from Fisher’s exact tests of associations between dome Up (C) or Down (D) DEGs and the developmental phases in A and B. The total number of each DEG expressed in each phase is indicated by the size of the point, and point fills indicate statistical significance. Dome Up or Down DEGs are over-represented in the Blastula/Gastrula or pre-ZGA phases, respectively. (E-F) Scaled QuantSeq counts for dome Up (E) or Down (F) DEGs across all developmental stages. Overall, dome Up or Down DEGs are normally up-or down-regulated by shield stage, so normal expression of dome DEGs is generally restored by shield. (G) Gene set enrichment analysis showing that genes known to be expressed during the first waves of zebrafish transcription are overexpressed upon Rif1 loss. The genes are ranked by Log2FC in expression between dome stage wild type and *rif1* omf201/omf201 embryos, as measured by QuantSeq. (H) Gene set enrichment analysis showing that genes, which are strictly zygotic or maternal, are often up-or down-regulated at dome stage in *rif1* mutants, respectively.

Using complementary approaches, Lee et al. and Heyn et al. defined “first wave” genes that are transcribed immediately upon zygotic genome activation **(*Heyn et al., 2014***; ***Lee et al., 2013***). We used gene set enrichment analysis to determine the extent to which the *rif1omf201/omf201* mutants overexpress first-wave genes (***Figure 6***G). At the dome stage, 272 out of 383 first wave genes had a Log2FC > 0 in mutants compared to wild-types. Furthermore, the first wave gene sets were located towards the top of the list of all genes ranked by the dome stage Log2FC (***Figure 6***G). Our data indicate that Rif1 loss generally increases the transcription of genes at the ZGA, but by including genes with substantial maternal contribution in our analysis, we might have underestimated that effect. Therefore, we used additional gene sets identified by Harvey et al. as being strictly zygotic or maternal (***Figure 6***H) **(*Harvey et al., 2013*)**. Like the “first wave” genes, the majority of genes in the zygotic set were expressed at higher levels (72/98 Log2FC > 0) in mutant embryos at the dome stage compared to wild-type, and most of the zygotic genes were towards the top of the list of genes ranked by dome Log2FC (***Figure 6***H top). In contrast, genes in the maternal set were often down in the mutants at dome and were found towards the bottom of the ranked gene list (***Figure 6***H bottom). Plotting mRNA levels for all Dome DEGs showed that Dome Up-DEGs are normally upregulated from pre-MBT to Shield stages, but exhibit earlier upregulation in *rif1* mutant embryos (***Figure 6***E). Conversely, Dome Down-DEGs normally decrease in expression between Dome and Shield stages, but show earlier reduction in mutants (***Figure 6***F).

Altogether our data suggest that Rif1 loss enhances the destruction of maternal mRNAs and stimulates the expression of genes during zygotic genome activation.

To directly assess transcription during ZGA, we labeled nascent mRNAs in early embryos with 4-Thiouridine (S4U) **(*Neumann et al., 2019*)**. By injecting the label into one-cell stage embryos we were able to identify mRNAs synthesized by the pre-MBT (2.5 hours) or dome (4.3 hours) stages. The incorporated label was mapped to mRNAs using thiol(SH)-linked alkylation for metabolic sequencing of RNA (SLAM-seq). SLAM-seq introduces T-to-C conversions at the sites of the incorporated S4U. Thus, each gene’s percentage of T-to-C conversions indicates the amount of its mRNA synthesized during the labeling period (zygotic) relative to pre-existing mRNA (maternal).

At the pre-MBT stage, there was very little difference in T-to-C conversion percentages between wild-type and *rif1omf201/omf201* mutant embryos, and there were only 102 differentially labeled genes (DLGs) at preMBT (***Figure 7***A-C; fold-change in T-to-C conversion > 1.5 and beta-binomial test FDR > 0.1). Therefore, we conclude that zygotic genome activation is not occurring earlier than the MBT for most genes in the *rif1omf201/omf201* mutants. In contrast, there were 1,872 DLGs with increased labeling in mutants at dome (***Figure 7***D, E; fold-change in T-to-C conversion > 1.5 and beta-binomial test FDR > 0.1). Furthermore, the T-to-C conversion rates at the dome stage were higher in the *rif1omf201/omf201* mutants for all gene categories analyzed (***Figure 7***F). The differences in T-to-C conversion were greatest for the genes we previously identified as being overexpressed in the mutants (256-Cell and dome Up-DEGs) and for the genes known to be transcribed during ZGA (First Wave and Zygotic). Notably, we detected a significant difference in mean T-to-C conversion calculated for all labeled genes. Overall, these data demonstrate that Rif1 has pervasive effects on transcription during zygotic genome activation.

**Figure 7.**
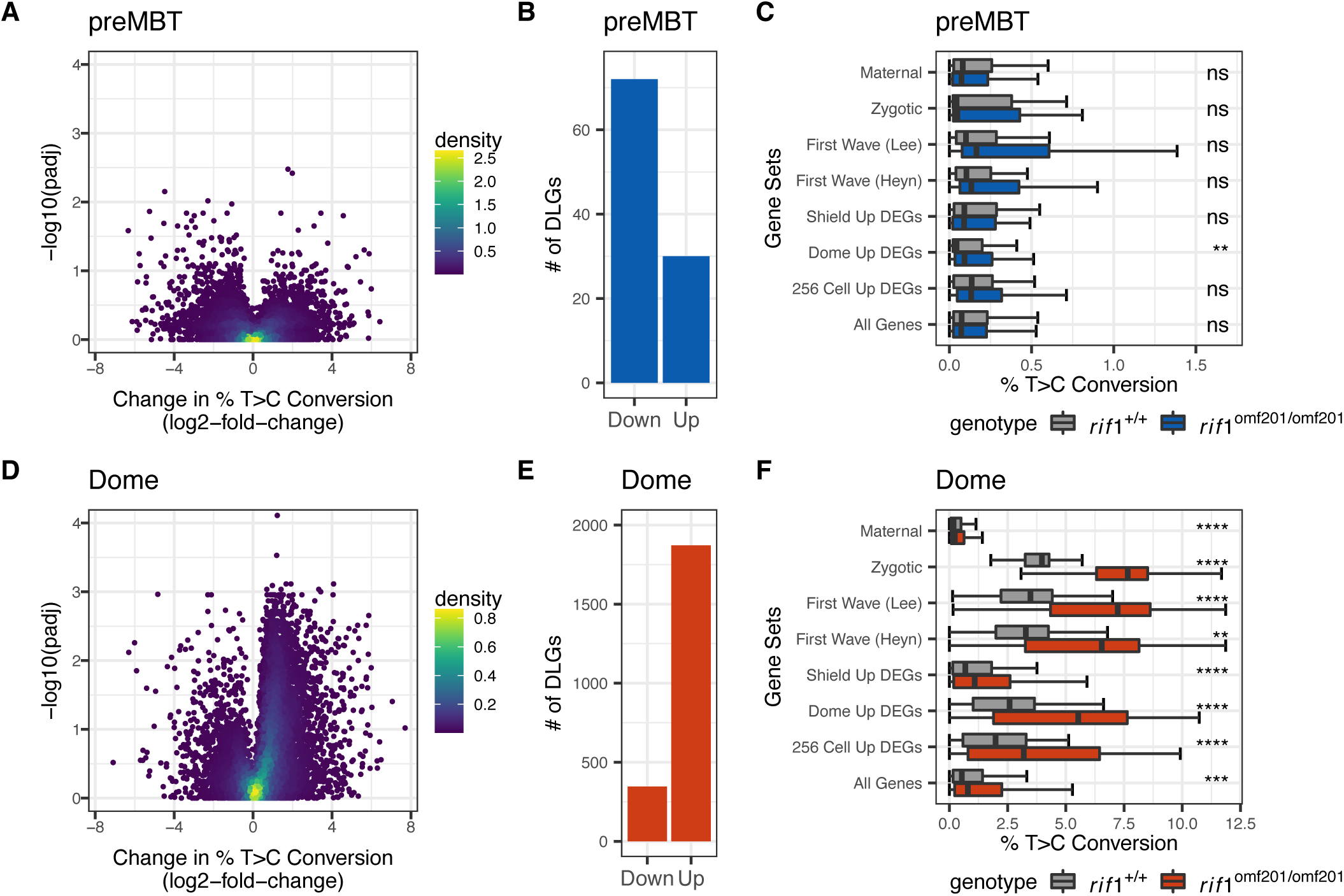
Labeling of nascent mRNAs reveals that the role for Rif1 in regulating ZGA transcription is pervasive. (A, D) Volcano plots showing the Log2-fold change in T-to-C conversions from SLAM-seq for each transcript between wild type and *rif1* mutant embryos at preMBT (A) or dome (D) stage. P values were calculated using the beta-binomial test and were corrected using the Benjamini & Hochberg method. The colors represent the density of overlapping data points. (B, E). Barplots of the number of differentially labeled genes (DLGs; >1.5-fold change in T-to-C conversions in *rif1omf201/omf201* mutants relative to wild-types and BH-adjusted p value <= 0.1) in A or D. (C, F) Boxplots of T-to-C conversion percentages at preMBT (C) or dome (F) stage for the following : all genes; DEGs upregulated in *rif1* mutants from this study (256 Cell Up, Dome Up, or Shield Up), genes transcribed during the first ZGA wave from Heyn et al. or Lee et al., or genes identified by Harvey et al. as maternal or zygotic **(*Harvey et al., 2013***; ***Heyn et al., 2014***; ***Lee et al., 2013***).

## Discussion

Previous studies of Rif1 in vertebrates have mainly been limited to cell culture models and have not addressed how its loss impacts RT establishment and change in the early embryo. We have developed a zebrafish model to study the developmental effects of Rif1 loss. Using this model, we have determined the impacts of Rif1 knockout on zebrafish embryonic development and survival, RT, and gene expression.

Our work adds to our understanding of the importance of Rif1 for vertebrate development. The extent to which Rif1 is essential for embryogenesis has been unclear. In *Drosophila melanogaster*, Rif1 knockdown by RNA interference caused embryonic lethality, but its knockout by CRISPR-Cas9 caused only mild effects on hatching rates(***Armstrong et al., 2020***; ***Seller and O’Farrell, 2018***; ***Sreesankar et al., 2015***). Rif1 knockout phenotypes in mice depend on genetic background and sex. Rif1 is essential in female mice, which die between the blastula and early gastrula stages(***Enervald et al., 2021***). Male mice are less affected by Rif1 loss. Although most Rif1 mutant male mice die around mid-gestation, a fraction survives to sexual maturity **(*Buonomo et al., 2009***; ***Chapman et al., 2013***; ***Daxinger et al., 2013***; ***Enervald et al., 2021***). We observed only mild *rif1* phenotypes during embryonic development. Rif1 loss only slightly delayed epiboly completion. The reason for that delay is unknown, but it may be due to the transcriptional effects of Rif1 loss at the ZGA. The *rif1* mutants also showed a deficiency in primitive hematopoiesis. The first wave of hematopoiesis is not essential for zebrafish development, as it is in mice. Thus, defects in hematopoiesis could possibly contribute to the poor survival of Rif1 mutant mice.

The most evident phenotype in the *rif1* mutant zebrafish is the substantial reduction in the number of females. It is tempting to speculate that this phenotype is related to female-specific lethality in mice, but we suspect the underlying causes differ. In part, Rif1 mutant female mice die due to an X chromosome inactivation defect, but zebrafish lack sex chromosomes **(*Enervald et al., 2021*)**. Furthermore, since we showed that expected numbers of *rif1* mutant fish survive to sexual maturity, the skew in gender ratio must be due to the preferential development of male fish rather than female lethality. A reduction in the proliferation or survival of primordial germ cells (PGCs) causes transdifferentiation of zebrafish to males **(*Tzung et al., 2015*)**. To understand how Rif1 promotes female development, we must determine whether its loss affects PGC proliferation. Intriguingly, mouse PGCs exhibit increased replication-transcription conflicts and are sensitive to mutations in Fanconi anemia pathway genes required for replication stress survival **(*Luo et al., 2014***; ***Yang et al., 2022*)**. Zebrafish PGCs exhibit high levels of intergenic transcription, which suggests that they may also have increased replication-transcription conflicts **(*Redl et al., 2021*)**. Furthermore, mutations in the Fanconi anemia DNA repair pathway disrupt female zebrafish development **(*Ramanagoudr-Bhojappa et al., 2018***; ***Rodríguez-Marí et al., 2010*)**. Therefore, the female development defect in the Rif1 mutant zebrafish could be due to deregulated replication fork initiation or impaired repli-cation fork stability in PGCs.

Studies with flies and mammalian cells have shown situations where Rif1 becomes less critical for RT once constitutive heterochromatin forms. For example, *D. melanogaster* Rif1 delays the repli-cation of specific satellite repeats before H3K9me3 and HP1a associate with them, but once constitutive heterochromatin forms, Rif1 is unnecessary **(*Seller and O’Farrell, 2018***; ***Yuan and O’Farrell, 2016*)**. In human HCT116 cells, late replication of genomic domains with high levels of the heterochromatin mark H3K9me3 does not depend on Rif1 **(*Klein et al., 2021*)**. In contrast, Rif1 knockout nearly completely ablates the RT pattern in human embryonic stem cells, which have relatively little constitutive heterochromatin **(*Hawkins et al., 2010***; ***Klein et al., 2021*)**. Pre-gastrula zebrafish embryos express high levels of Rif1 and substantially lack constitutive heterochromatin (***Figure 1***C and **(*Laue et al., 2019*)**). Thus, we were surprised that Rif1 loss had relatively little effect on RT in early embryos. Instead, we found that RT differences were more pronounced later in development. Understanding this paradox will likely reveal mechanisms that control Rif1 activity.

Although the effects of Rif1 loss on RT increased as zebrafish development proceeded, the significant developmental changes that shaped the RT profile of late-stage embryos were not affected. For example, *rif1* was not required for the early-to-late timing switches that occur over large segments of chromosomes 4 and 22 during gastrulation, nor was it required for the widespread late-to-early switches that coincided with enhancer activation. Much of the variance in RT between wildtype and mutant embryos was due to a general flattening of the timing profiles, and Rif1 loss did not disrupt the overall peak and valley profile structure. Therefore, our data suggest that instead of causing developmental RT switches in zebrafish, Rif1 sharpens RT caused by other unknown determinants.

If Rif1 is not required, what drives developmental changes in RT? Zebrafish embryos establish constitutive heterochromatin by the start of gastrulation, so the early-to-late RT switches that occur during and after that stage might be due to further maturation of heterochromatin **(*Laue et al., 2019*)**. The activation of enhancers might drive late-to-early RT changes throughout the zebrafish genome. Early replicating control elements (ERCEs), which have active epigenetic marks associated with enhancers, cause early replication of large domains in the mouse genome **(*Sima et al., 2019*)**. More work is needed to determine whether the formation of constitutive heterochromatin and activation of ERCEs drive developmental changes in RT in zebrafish.

We were particularly interested in whether RT changes in the embryo would lead to transcriptional changes. A long-standing hypothesis is that the RT of a gene can affect its transcriptional competence **(*Gottesfeld and Bloomer, 1982***; ***Lande-Diner et al., 2009***; ***Zhang et al., 2002***). Recently, Klein et al. used human cell lines in which Rif1 could be conditionally depleted to show that its loss caused widespread and coordinated RT and epigenomic changes **(*Klein et al., 2021*)**. Furthermore, the epigenomic alterations caused by Rif1 loss were often associated with gene expression changes **(*Klein et al., 2021*)**. Although this work strongly supports the hypothesis that RT main-tains the epigenome, the effects of Rif1 loss in a developing embryo where the RT program and the epigenome are established have not been examined. We used whole-genome RT and RNAseq analysis to determine whether Rif1 loss caused RT changes that led to deregulated gene expression. Unexpectedly, we found that when Rif1 had the most significant effect on RT during later development, there were few changes in mRNA levels. One limitation of the current work was that we did not directly measure alterations in the epigenome, which may have occurred without transcriptional changes. A second limitation was that we measured RT and gene expression for all cells in the embryo in bulk. Consequently, we were unable to detect cell-to-cell variability in either RT or gene expression. Indeed, RT deregulation could cause epigenetic variability. Nonetheless, our data show that RT changes caused by Rif1 loss do not substantially change overall gene expression later in development.

A further limitation of this study is that we concentrated on replication timing without directly measuring other features of the replication program that contribute to this timing. These features include origin usage, replication fork spacing, fork directionality, fork progression, and fork stability. A more comprehensive understanding of how Rif1 loss affects these parameters will be important for defining the relationship between Rif1-dependent changes in replication timing and transcription.

The most pronounced effects of Rif1 loss on transcription were during the earliest stages of development. We previously showed that RT patterns are detected around the MBT in zebrafish embryos, but those patterns are relatively weak compared to later development. In this work, we showed that there was almost no detectable RT pattern in wild-type or Rif1 mutants before the ZGA. Despite the lack of RT patterns, Rif1 loss affected the first major wave of transcription. Most genes affected by Rif1 loss at that stage increased slightly, but a significant fraction of all genes transcribed at the ZGA were upregulated. Thus, Rif1 loss had a strong overall effect on transcription at the ZGA. Notably, the role for Rif1 in regulating ZGA transcription is evolutionarily conserved. As we show here for zebrafish, mouse pluripotent cells express Rif1 at high levels, and Rif1 expression decreases during cellular differentiation **(*Adams and McLaren, 2004*)**. Multiple groups have reported that Rif1 loss in mouse embryonic stem cells (mESCs) causes increased expression of twocell (2C) zygote-specific genes **(*Dan et al., 2014***; ***Li et al., 2022***, ***2017b***; ***Yoshizawa-Sugata et al., 2021***; ***Zhang et al., 2022*)**. Given that ZGA occurs through different transcription factors in zebrafish and mice, it is likely that Rif1 affects genes expressed at the ZGA in both species by regulating a common fundamental process rather than the activities of specific transcription factors. mESCs in the 2C state have increased histone marks associated with active transcription, and Rif1 loss is sufficient to cause these chromatin changes **(*Li et al., 2017a***; ***Yoshizawa-Sugata et al., 2021***). Mouse Rif1 co-immunoprecipitates with multiple repressive histone methyltransferases and with the noncanonical polycomb repressive complex PRC1.6, and Rif1 promotes the recruitment of chromatin regulators to several 2C genes in mESCs **(*Li et al., 2017a*)**. We do not know how Rif1 affects the epigenome in the early zebrafish, and this is an important goal for future research. Given the striking similarity in Rif1 function in early zebrafish and mouse embryos, we anticipate the role of Rif1 in regulating transcription in the early embryo will be broadly conserved in vertebrates.

A recent study in mouse embryos independently identified RIF1 as a regulator of the developmental consolidation of the RT program. RIF1 depletion produced a less-defined, developmentally immature RT program, while RIF1-dependent RT changes were not correlated with transcriptional changes **(*Nakatani et al., 2025*)**. Together with our findings in zebrafish, these results support a conserved role for RIF1 in sharpening replication timing during vertebrate development and indicate that its effects on replication timing can be uncoupled from changes in gene expression.

In conclusion, we have described multiple phenotypic consequences of Rif1 loss in zebrafish. We found that Rif1 undergoes a functional switch during zebrafish development. When it is highly abundant during early development, Rif1 regulates transcription but does not affect RT. Later in development, Rif1 is important for RT but does not substantially affect gene expression. The zebrafish will be useful for understanding how Rif1 switches between these functional modes at different stages of development. Furthermore, we have identified a new function for Rif1 in sex determination. Thus, the *rif1* mutant zebrafish described here will serve as a valuable model for understanding PGC biology and sex determination.

## Methods and Materials

### Animal care

All animals were handled in strict accordance with protocols approved by the OMRF Institutional Animal Care and Use Committee. Adult breeding fish were housed in an aquatic animal facility in tanks of ∼30 fish per 3 liter tank and maintained at 26.5◦C with 10-hour light and 14-hour dark cycles.

### TALEN production and microinjection

The TALEN expression constructs targeting the seventh exon of the *rif1* gene were assembled in pCS2TAL3-DD and pCS2TAL3-RR using Golden Gate Assembly as in Cermak et al. **(*Cermak et al., 2011*)**. pCS2TAL3-DD and pCS2TAL3-RR were gifts from David Grunwald (Addgene plasmids # 37275 and 37276; RRID:Addgene_37275; RRID:Addgene_37276). mRNA encoding the TALE nuclease subunits was synthesized using the mMESSAGE mMACHINE SP6 Transcription Kit (AM1340; ThermoFisher Scientific, Waltham, MA) and purified using RNeasy Mini Kit (74104; Qiagen). 125pg of each TALEN mRNA was injected into 1-cell stage zebrafish embryos. TALEN-injected embryos were raised to adulthood and assessed for transmission of insertions/deletions at the TALEN cut site using high resolution melt analysis (HRMA). F1 fish from mutation transmitting F0 animals were raised to adulthood and analyzed for mutation by HRMA, and the specific *rif1* mutation was identified by Sanger sequencing of PCR amplicons spanning the mutation site. *rif1* mutant fish were genotyped using HRMA, Sanger sequencing, and/or competitive allele-specific PCR (KASP assay; LGC).

### Zebrafish embryo collection and processing for gDNA, mRNA, or protein

Zebrafish were spawned as described previously **(*Siefert et al., 2018*)**. For SlamSeq, 2-cell embryos were injected with 50 pmoles of 4-Thiouridine (S4U). Embryos were incubated at 28.5◦C. Before collection, clutches were visually inspected to ensure that all embryos were developing synchronously. Unfertilized or morphologically abnormal embryos were removed before collection. Whole embryo pools to be used for gDNA or mRNA were rapidly frozen and stored at -80◦C for later processing. Embryos were prepared for fluorescence activated cell sorting based on DNA content as described in Siefert et al(***Siefert et al., 2018***). Briefly, bud stage or 24 hpf embryos were dechorionated with Pronase, deyolked, and then triturated to a single cell suspension in Phosphate Buffered Saline. Nuclei were prepared from the cells were incubated in DNA staining solution (PBS; 1% Bovine Serum Albumin; 50 ug/ml Propidium iodide P4170-25mg Sigma-Aldrich; 100 ug/ml RNAse A 12091039 Invitrogen) on ice for at least one hour before sorting. The nuclei were then filtered through a 40*μ*m mesh before analysis on a BD LSRII (shield and bud), BD Biosciences FACSAria IIIu (28hpf) or BD Biosciences FACSCalibur. Genomic DNA was prepared from whole embryo pellets or sorted nuclei using the Quick-DNA Miniprep Plus Kit (D4069; Zymo Research). mRNA was prepared from whole embryos using the RNeasy Mini Kit (74104; Qiagen). For protein lysates, whole embryos in pools were deyolked and disaggregated into a single cell suspension as previously described **(*Link et al., 2006*)**, and dry cell pellets were stored at -80◦C for later use. To make crude nuclear extract, cell pellets were resuspended in Buffer A (10mM HEPES, 10mM KCl, 1.5mM MgCl_2_, 0.34M sucrose, 10% glycerol) with 0.1% triton and protease inhibitors followed by Buffer A plus inhibitors, 0.1% triton and 1uL/mL Benzonase (E1014; Sigma).

### *In situ* hybridization

The Thisse Lab protocol was followed for the ISH **(*Thisse and Thisse, 2008*)**. Briefly, ISH probes were constructed by PCR from genomic DNA and isothermal assembly into HindIII-BamHI digested pUC57-Amp plasmids flanked by T3 and T7 promoters for sense (coding RNA) and anti-sense strand synthesis, respectively. Primers used to amplify probe sequences are as follows: *b-globin* forward acc ctc act aaa ggg aaC GCG CAG CGA TTC AGA ACA T, *b-globin* reverse acg act cac tat agg gcC TTC TGA GGG CTG ACA CAA CA, *gata1a* forward acc ctc act aaa ggg aaA CTC CTC TGA GCC TTC TCG T, and *gata1a* reverse acg act cac tat agg gcT GTG GAG AAG GGC CGT AAA C. Dioxygenin (DIG)-labelled RNA probes were synthesized with T3 and T7 polymerase with reagents from Roche. Embryos were dechorionated with forceps and fixed in 4% paraformaldehyde in 1X PBS overnight. Fixed embryos were dehydrated in 100% methanol and stored at -20◦C. The 24 hpf embryos were proteinase K treated for 10 minutes. The protocol was programmed into an Intavis VSi machine to automate the prehybridization and washing procedures. Embryos were removed from the machine and incubated in DIG-labelled RNA probes overnight in a 70◦C water bath. Embryos were then returned to the machine for washing, preincubation with Bovine Serum Albumin, incubation with anti-DIG antibody, and additional washing. Labelled embryos were imaged with a Nikon SMZ1500 stereomicroscope with Andor Zyla sCMOS camera.

### Antibodies used for immunoblotting

Anti-zebrafish Rif1 antibodies were made by immunizing rabbits with bacterially expressed purified fragments of zebrafish Rif1 (NP_001074275.1 amino acids 1-150) and were affinity purified from whole serum (Cocalico Biologicals, Inc.). Rabbit polyclonal anti-human Smc3 was a gift from Susan-nah Rankin **(*Lafont et al., 2010*)**. Peroxidase AffiniPure Goat Anti-Rabbit IgG (H+L) was purchased from Jackson Immuno Research Laboratories (RRID: AB_2313567).

### Quantitative RT-PCR

mRNA was converted to cDNA using the AccuScript High Fidelity Reverse Transcriptase (600089; Agilent, Santa Clara, CA). qPCR was done on a LightCycler 480 with the SYBR Green I Master (04707516001; Roche) master mix. Three technical replicates were performed for each sample. The relative mRNA expression was calculated using the 2-ΔΔCq method. The following primer sequences were used: *rif1* forward ACT GGC TGA TGA CAT TGA TCG T, reverse ATC CAG GGC TAC GGA GGT TA; *eef1a1l1* forward CTG GAG GCC AGC TCA AAC AT, reverse ATC AAG AAG AGT AGT ACC GCT AGC ATT AC.

### Next generation library preparation and sequencing

For whole-genome sequencing, 50ng - 2*_μ_*g of gDNA was sheared to approximately 300 bp fragments using a sonicator (Covaris). The gDNA libraries were prepared using the KAPA Hyper Prep Kit (Roche) with SeqCap (Roche) or Illumina TruSeq (Illumina) adapters. Genomic DNA libraries were sequenced in 150bp paired-end read fragments on an Illumina HiSeq 2500 or NovaSeq 6000. Before library preparation, the SlamSeq mRNA was Iodoacetamide treated using the SLAMseq Kinetics Kit – Anabolic Kinetics Module (Lexogen). The QuantSeq and SlamSeq libraries were prepared using the QuantSeq 3’ mRNA-Seq Library Prep Kit for Illumina (Lexogen). RNA sequencing was performed using custom primers on an Illumina Nextseq 500 with High Output chemistry and 75bp single-ended reads.

### Replication timing analysis

RT data were generated based on copy-number variations between filtered G1 and S-phase wholegenome sequencing read counts, as described previously (Siefert, 2017). Pools of >100 whole embryos at preMBT, dome, or shield stage were treated as S-phase samples, since a high percentage cells at these stages are in S phase. Nuclei from bud or 24 hpf embryos were sorted using a BD Biosciences FACSAria IIIu based on DNA content to isolate G1 and S-phase fractions. The data was processed using our RepliTimer pipeline (DOI 10.5281/zenodo.6428748) **(*Sansam, 2022a*)**. Whole genome sequencing reads were trimmed with fastp, aligned to the zebrafish genome (GRCz11) using BWA-MEM, and duplicate reads were marked with Picard MarkDuplicates. Properly-paired and non-duplicated reads with MapQ scores of at least 20 were filtered with Bamtools. Alignments were read into GenomicRanges for downstream analysis in R. Reads were also removed if they were within a set of 100 bp windows with anomalously high read counts (exceeding 1.5 times the interquartile range in at least 4 out of 65 whole genome sequencing runs from this study and previously published work) **(*Siefert et al., 2017*)**. Genomic windows for calculating RT were defined as regions with a minimum of 400 reads in every sample. An S/G1 quotient was calculated for each window, and quotients were smoothed and Log2 transformed **(*Siefert et al., 2017*)**.

### RNA sequencing analysis (QuantSeq 3’ mRNA-Seq and SLAM-Seq)

The QuantSeq data was analyzed using our QuantSeqPipeline pipeline (DOI 10.5281/zenodo.6622165) **(*Sansam, 2022b*)**. Reads were trimmed with bbmap and aligned to GRCz11 with STAR. Using a custom R script, probable internal priming sites at polyA genomic sequences were identified and reads mapping to those sites were removed. Reads aligning to 3’ ends of transcripts are then counted. The 4.3.2 transcript annotation by Lawson et al was used **(*Lawson et al., 2020*)**. Differentially expressed genes were identified using DESeq2, and plots were generated in R. The SLAM-Seq sequencing reads were aligned and T-to-C conversion rates were calculated using the SLAM-DUNK package with default settings **(*Neumann et al., 2019*)**.

## Availability of data and materials

The data discussed in this publication have been deposited in NCBI’s Gene Expression Omnibus and are accessible through GEO Series accession numbers GSE225957, GSE226033, and GSE226222. The plasmids pCS2TAL3-RR-zrif1-exon8 (Addgene plasmid 194434), pCS2TAL3-DD-zrif1-exon8 (Adbodgene plasmid 194435), pUC57-GATA1 (Addgene plasmid 194436), and pUC57-b-globin (Addgene plasmid 194437) have been deposited at Addgene.

## Acknowledgements

We thank the Imaging, Flow Cytometry, Clinical Genomics, and Research Computing Core Facilities at the Oklahoma Medical Research Foundation for their assistance with microscopy, cell sorting, sequencing, and computing, respectively.

## Ethics approval and consent to participate

All animals were handled in strict accordance with protocols approved by the OMRF Institutional Animal Care and Use Committee.

---

**Figure 5—figure supplement 1.**
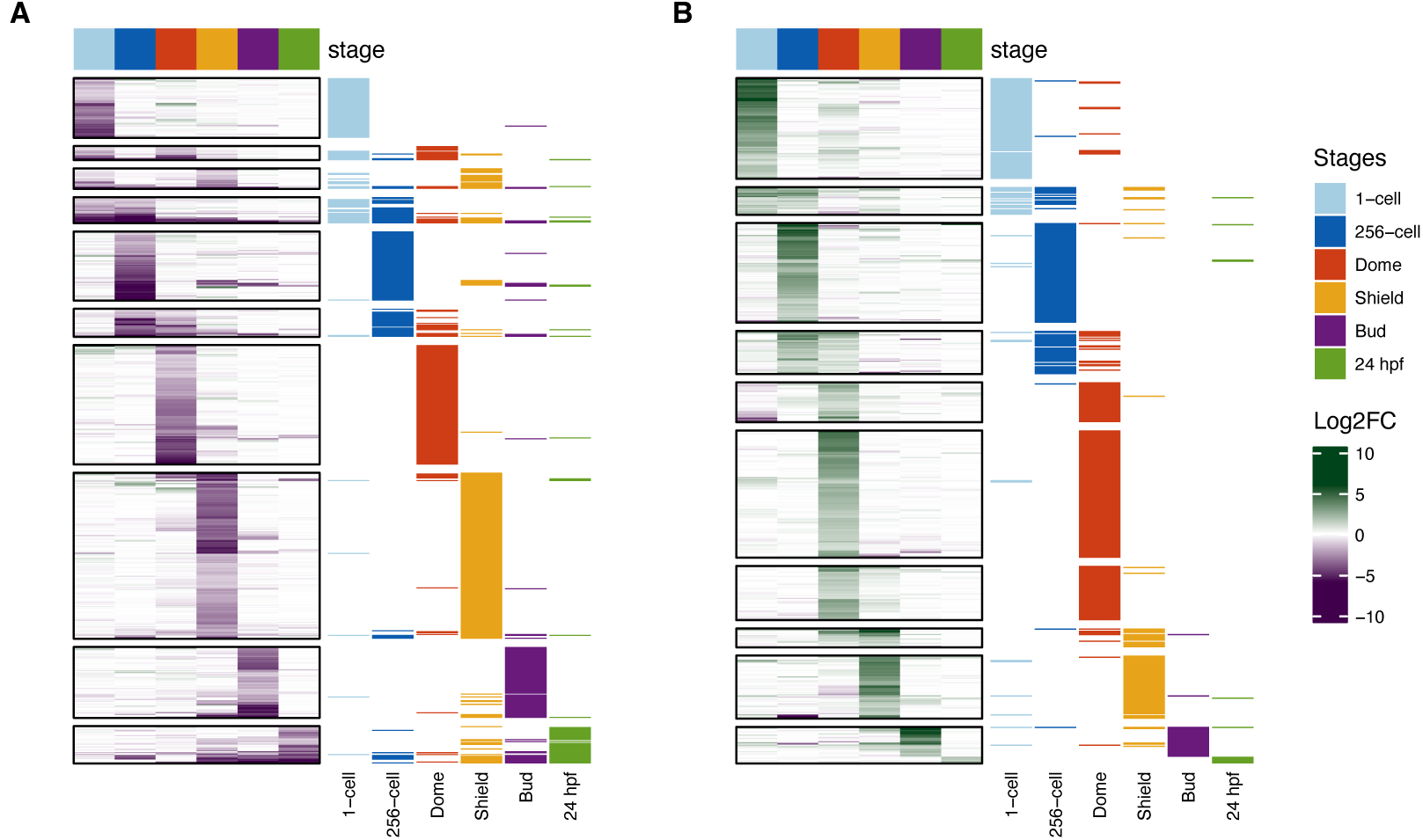
Most DEGs are at normal levels in maternally-contributed mRNA. Heat plots showing the Log2 fold change in mRNA expression of *rif1* mutants vs wild types for all Down-DEGs (A) or Up-DEGs (B). Genes are displayed in rows and developmental stages are in columns. Rows were split by k-means clustering. Colors in the row annotations to the right of each heat plot denote the stage(s) at which each DEG was identified. The majority of 1-cell DEGs are expressed normally at other stages, and the majority of DEGs at other stages are expressed normally at 1-cell.

**Figure 5—figure supplement 2.**
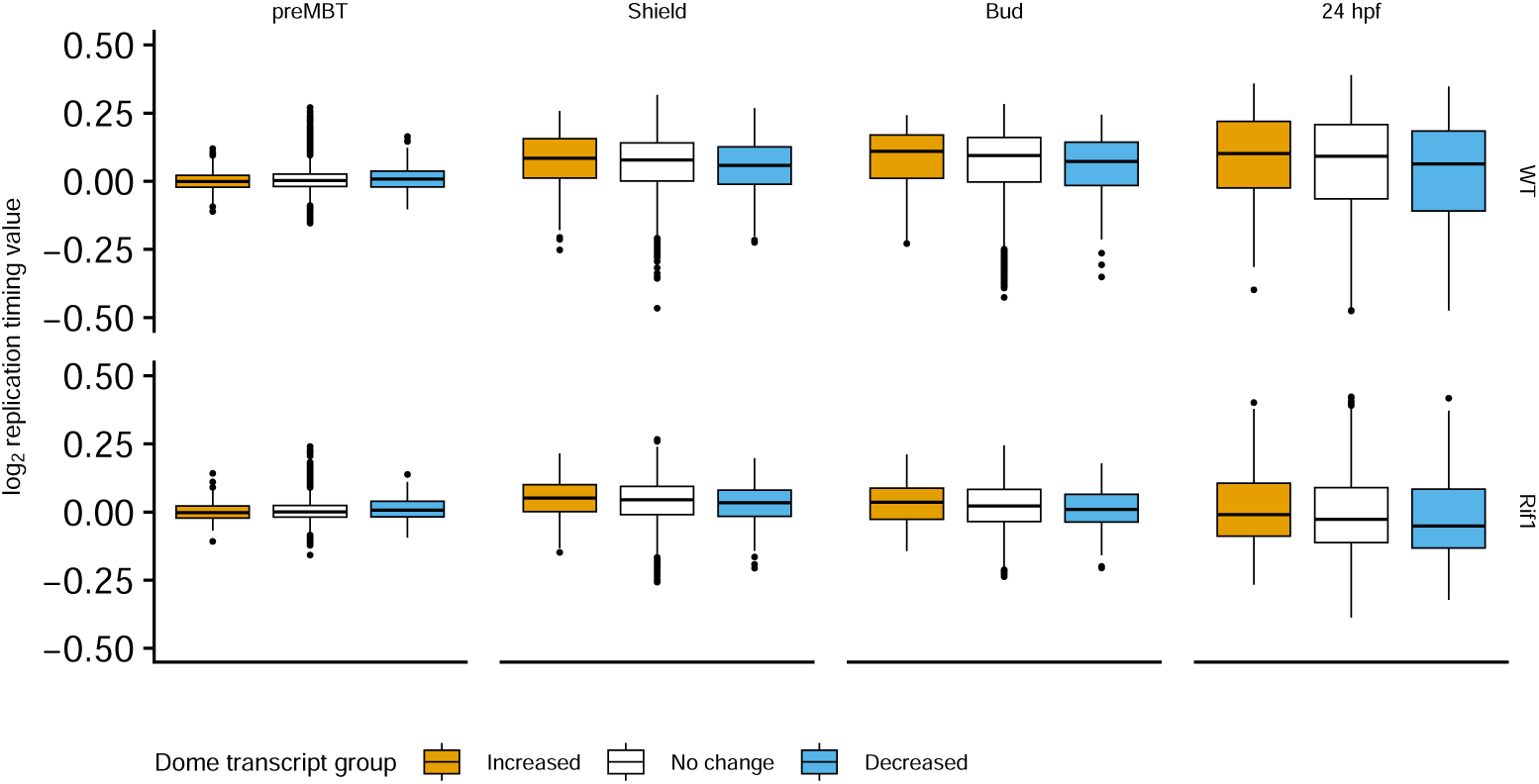
Replication timing values of genes with altered transcript abundance at Dome. Boxplots show log2-transformed smoothed replication timing values assigned to genes by the nearest replication timing bin. Genes were grouped according to differential transcript abundance in *rif1* mutant embryos at Dome: increased, decreased, or not significantly changed. Replication timing values are shown across developmental stages and genotypes. Genes with increased or decreased transcript abundance at Dome did not show a clear enrichment for early- or late-replicating regions relative to genes with no significant transcript change.

